# Spatial Development of Brain Networks During The First Six Postnatal Months

**DOI:** 10.1101/2024.09.19.613960

**Authors:** Masoud Seraji, Sarah Shultz, Qiang Li, Zening Fu, Vince D Calhoun, Armin Iraji

## Abstract

The initial months of life constitute a crucial period for human development. A comprehensive understanding of this early phase is essential for unraveling the origins of neurodevelopmental disorders and promoting infant brain health. This study uniquely focuses on the spatial development of intrinsic brain connectivity networks during infancy, which has been less explored compared to functional connectivity. We utilized independent component analysis on resting-state fMRI data from 74 infants to assess how the spatial organization of infant brain networks evolves between birth and six months.

Our findings reveal significant changes in spatial characteristics, including an a notable rise in the network-averaged spatial similarity across age, reflecting how closely each participant-specific spatial map aligns with the group-level map for each network. We also observed a marked reduction in the network engagement range by age, representing the extent of voxel intensity range fluctuation within each network. This suggests a continuing process of consolidation, where voxel contributions to the network become more uniform, as indicated by the narrowing of intensity values. The network strength, calculated as the average of all the voxel intensities in the network, indicating the degree of involvement to the specific functional network, increased across age in several networks, such as frontal-mPFC, primary, and secondary visual networks. The network size, along with the network center of mass, illustrating spatial distribution alterations of brain networks by age, varied across different networks. For instance, both metrics increased across age in the secondary visual network but decreased in the temporal network. Additionally, we examined the networks in relation to their linear versus non-linear developmental trajectories across all spatial characteristics, providing a deeper understanding of how these patterns evolve during early infancy. These findings contribute to early brain development understanding and offer insights into potential markers of consolidation and spatial reorganization in large-scale brain networks during infancy.

## 1. Introduction

The first few months after birth are considered a crucial timeframe in which the fundamental aspects of the brain’s functional and structural framework are established [1,2]. Consequently, there is growing interest in understanding how deviations from typical neurodevelopment during this period may play a pivotal role in the emergence of developmental disorders and psychopathology [3]. Resting state functional connectivity (FC)— the measurement of correlated spontaneous low-frequency fluctuations in the blood-oxygen-level dependent fMRI signal across functionally related brain regions during rest—provides a window into functional brain organization in early infancy [4,5]. It is well-established that most adult functional brain networks are observable from birth (perhaps even in fetal period) and undergo asynchronous developmental changes over first postnatal months and years [6–11]. More recently, studies have begun to identify deviations in these critical periods of FNC development that are associated with the emergence of developmental disability, further highlighting the value of charting trajectories of functional brain network maturation.

While existing studies of FNC in infancy have shed light on developmental change in the temporal coupling within and between networks, developmental change in the spatial properties of functional connectivity patterns remains unexamined. Brains are known to be spatially dynamic (that is, the spatial configuration of functional regions change over time), and spatially dynamic analyses have revealed critical information about brain functional architecture and its association with behavior [12] and alterations in conditions such as schizophrenia [13] providing new insights into functional brain networks that may be hidden by existing approaches that focus on variations of temporal copuling among spatially fixed functional sources [13].

Here, we provide the first investigation of developmental changes in spatial organization of functional brain networks during infants’ first 6 postnatal months, a period characterized by extremely rapid brain growth and development [14,15]. Infants completed up to 3 scans during their first 6 postnatal months, providing dense longitudinal coverage over this highly dynamic period of development. While our methodology closely aligns with many studies employing ICA on fMRI data in adults, which typically involves group-level ICA followed by back-reconstruction [16], we have enhanced our back-reconstruction process compared to previous infant studies. This improvement involves the adoption of group information-guided ICA (GIG-ICA) [17], which enhances spatial correspondence across participants. As a result, we achieve improved spatial accuracy when estimating participant-specific spatial maps [17]. We mapped trajectories of change in several measures of spatial characteristics—network-averaged spatial similarity, network engagement range, network strength, network size, and network center of mass—to understand how functional networks change during early infancy. Network-averaged spatial similarity provides insights into the consistency of network configurations among individuals, allowing researchers to assess how well a participant’s brain network matches commonly shared spatial patterns. Network engagement range helps identify how consistently or variably brain regions are involved within the network. Network strength revealing critical insights into how strongly different brain regions are participating in the network’s function, while network size provides a measure of brain growth or pruning processes, key to understanding network expansion or contraction in response to development. Lastly, network center of mass offers clues about the spatial organization and balance of brain functions. Overall, critical examination of spatial characteristics is essential due to the considerable spatial variability observed among individuals. This highlights the need for adaptive analytical approaches that can accurately capture the unique functional regions in each participant [12]. Overlooking these spatial differences risks misinterpreting the underlying functional data, leading to inaccurate conclusions about brain activity and network configurations [12]. Therefore, incorporating spatial measures allows for a more precise understanding of brain function and developmental trajectories.

In line with the rapid brain changes unfolding during this period [18], we predicted that spatial maps would undergo significant change during infants’ first postnatal months. Specifically, given that FNC development generally follows a sequence from primary to higher-order networks—with primary networks showing significant changes within the first three postnatal months, and higher-order networks exhibiting fewer changes in the first six postnatal months [9]—We hypothesized that the spatial maps of both primary and higher-order networks would experience changes from birth to six months. By mapping longitudinal change in the spatial organization of functional networks, this study offers insight into a critical aspect of functional brain network maturation that was previously overlooked in studies of early infancy.

## 2. Materials and Methods

### 2.1. Participants

Participants were n=74 neurotypical infants (43 males and 31 females) enrolled in prospective longitudinal studies at the Marcus Autism Center in Atlanta, GA, USA (see Table 1 for participant demographics). Infants had a mean gestational age at birth of 39.2 weeks (SD=1.34) and were considered neurotypical on the basis of having no family history of autism in up to third-degree relatives, no developmental delays in first-degree relatives, no pre- or perinatal complications, no history of seizures, no known medical conditions or genetic disorders, and no hearing loss or visual impairment. Infants with contraindications for MRI were excluded.

**Table 1:** Demographics of participant sample. For information categories with a participant number less than the total sample, the N is specified next to the category title.

| **(N = 74)** |  |
| --- | --- |
| **Infant Sex** | 31f, 43m |
| **Gestational Age at Birth, mean (SD) (N=74)** | 39.2wks (1.34) |
| **Race (N=66)** |  |
| Black | 7.6% |
| White | 86.3% |
| More than one race | 6.1% |
| **Maternal Education (N=64)** |  |
| High School | 1.6% |
| Trade/Vocational Training | 1.6% |
| College Courses | 6.2% |
| Associate's Degree | 1.6% |
| College Degree | 34.3% |
| Graduate Degree | 54.7% |
| **Household Income (N=63)** |  |
| < \$40000 | 4.8% |
| \$40000-\$80000 | 19.0% |
| \$80001-\$100000 | 22.2% |
| \$100001-\$150000 | 28.6% |
| > \$150001 | 25.4% |

Participants’ parents provided informed written consent, and the Emory University Institutional Review Board approved the research protocol for this study.

Scans were scheduled for each infant at up to 3 pseudorandom timepoints between birth and 6 months (Fig. S1). This approach yielded dense coverage over the 0- to 6-month period (137 scans, each separated by a mean of only 1.4 days (SD=1.8); see Fig. S1 in section 1.2 of supplemental for a distribution of participants age at each scan), with 28, 29, and 17 of infants contributing one, two, or three longitudinal scans, respectively. Infant age at each scan was corrected for length of gestation, defined as the age in days minus 7*(40-gestational age in weeks) [19].

### 2.2. Data Collection

Infant scans were acquired at Emory University’s Center for Systems Imaging Core on a 3T Siemens Tim Trio (n=65) or a 3T Siemens Prisma (n=72) scanner, using a 32-channel head coil. All infants were scanned during natural sleep. Infants were swaddled, rocked, and/or fed to encourage natural sleep. Once asleep, infants were positioned on a pediatric scanner bed. To minimize scanner noise to levels below 80 dBA, two measures were implemented: (i) the use of sound attenuating pediatric headphones, featuring MR-safe optical microphones for real-time monitoring of in-ear sound levels, and (ii) the integration of a custom-built acoustic hood within the MRI bore [20].

To mitigate the abrupt onset of scanner noise, white noise through the headphones before commencing the first sequence. This white noise—gradually increased in volume—was played through the headphones prior to the first sequence. An MRI-compatible camera (MRC Systems) was affixed to the head coil, facilitating continuous monitoring of the infant during the scan. An experimenter was present in the scanner room throughout the scan and the procedure was stopped if the infant awoke or if an increase in sound level was observed.

Functional MRI data from the Tim Trio scanner were acquired using a multiband Echo planar imaging (EPI) sequence. The acquisition parameters were as follows: repetition time (TR) of 720 ms, echo time (TE) of 33 ms, flip angle of 53°, a multiband factor of 6, a field-of-view (FOV) of 208 × 208 mm, and an image matrix of 84 × 84. The spatial resolution was 2.5 mm isotropic, and 48 axial slices were acquired to cover the whole brain. To improve the signal-to-noise ratio (SNR), an additional 6 dummy scans were performed prior to the actual data acquisition to ensure steady-state magnetization.

Functional MRI data from the Prisma scanner were acquired using a multiband EPI sequence. The key acquisition parameters included a repetition time (TR) of 800 ms, an echo time (TE) of 37 ms, and a flip angle of 52°. The multiband factor was set to 8, with a FOV of 208 × 208 mm and an image matrix of 104 × 104. The spatial resolution achieved was 2 mm isotropic, encompassing 72 axial slices to cover the entire brain. To enhance the SNR, 6 dummy scans were performed before the actual data acquisition to ensure steady-state.

### 2.3. Preprocessing

Data preprocessing was conducted as follows: Initially, the first ten volumes, which exhibited significant signal changes for magnetization equilibrium purposes, were discarded. Head motion correction was performed using the mcflirt function in FSL (https://fsl.fmrib.ox.ac.uk/fsl/fslwiki/FSL), aligning all volumes to the first one. To correct for distortion in the multi-band rs-fMRI data, single-band data acquired with phase encoding in both the AP and PA directions were used to estimate the susceptibility-induced off-resonance field. Following distortion correction, slice timing correction was applied.

A two-step normalization process was employed to align the infant datasets into a common Montreal Neurological Institute (MNI) space. The first step involved normalizing the infant fMRI data to the 3-month UNC/UMN baby connectome project (BCP) T1 template [21]. In the second step, the output from the first normalization was further aligned to the common MNI space using the EPI template [22]. Finally, a 6 mm Gaussian kernel was applied to smooth the spatially normalized fMRI images.

### 2.4. Data Analysis

Quality control (QC) was conducted on the preprocessed data to ensure accuracy. Scans with poor normalization to the MNI space were excluded by comparing the individual masks with the group mask. Ultimately, no scans were excluded due to normalization issues or having fewer than 100 time points (see section 1.1 in the supplemental materials for detailed QC procedures). In addition, instead of excluding scans with significant head motion, we opted to retain all scans and account for head motion by including it as a covariate in our statistical models. This approach allows us to control for the influence of motion on our results, ensuring that any detected brain activity patterns are not confounded by movement artifacts (see section 2.7 for details of statistical analysis).

We conducted group ICA on the scans acquired from all infants [23], using the group ICA of fMRI toolbox (GIFT; http://trendscenter.org/software/gift) [24–26]. Initially, participant-specific principal components analysis (PCA) was utilized to standardize the data and ensure that each participant’s contribution is comparable within the shared subspace. This participant-specific PCA process also offers advantages in noise reduction and computational efficiency [27,28]. Thirty principal components (PCs) capturing the maximum variance for each participant were retained for subsequent analysis. Participant-specific PCA emphasizes individual differences at the participant level, whereas group-level PCA emphasizes commonalities among participants [28]. To implement this, all principal components derived from individual participants were combined along the time dimension, and then group-level PCA was performed on this concatenated dataset.

We utilized the top 20 group-level PCs explaining the maximum variance as input for group ICA. We applied a group ICA model order of 20, as suggested by previous studies [29,30], aimed to capture large-scale brain networks. To ensure result reliability, ICA was performed 100 times using the Infomax optimization algorithm [31] with random initialization and bootstrapping. The independent components most closely aligned with the centroids of stable clusters were deemed the optimal ones and chosen for subsequent analysis. We assessed the reliability and quality of the large-scale brain networks using the ICASSO quality index (IQ), quantifying component stability across runs [32]. In the fMRI large-scale brain networks estimation context, ICASSO IQ is commonly employed to distinguish reliable components suitable for further analysis from unstable ones [13]. A large-scale brain network was identified if it met specific criteria, including an ICASSO IQ value exceeding 0.80, high spatial overlap with gray matter, peak weight within gray matter, and low spatial similarity to motion, ventricular, and other artifact components.

Subsequently, we employed the GIG-ICA technique to estimate participant-specific large-scale brain networks [17]. After delineating the large-scale brain networks, we categorized each component by discerning their activation patterns within spatial maps. We ascribed labels to significant intrinsic connectivity networks (ICNs) based on their functional roles within the brain.

### 2.5. Age-Associated Voxel-Level Analysis

To focus on the regions within each brain network where developmental changes are most likely to occur, we explored the relationship between each network and corrected age through voxel-level correlation analyses. This approach involved assessing the correlation between voxel-specific activity in participant-specific spatial maps and corrected age across all scans. The results are presented as spatial maps, highlighting regions within each network where voxel activity shows a significant correlation with age.

After identifying voxels significantly associated with age within each network, we delved into exploring the interrelationship among these voxels across different networks. To conduct this analysis, we first computed the average intensity of all age-significant voxels within each network, a metric we refer to as the age-correlated voxel mean (ACVM). We then analyzed the correlations between these ACVMs across all networks, producing a 13 by 13 matrix illustrating the similarity of ACVMs between different networks, providing insight into how these values align across various brain networks. We next employed the modularity metric from the Brain Connectivity Toolbox (BCT) to quantify the extent to which the network could be segmented into distinct, well-defined groups (https://sites.google.com/site/bctnet/) [33]. It is important to note that for this specific analysis, we focused exclusively on the voxels that were significantly associated with age. However, for the remainder of the methodology, we conducted our analysis across all voxels within the identified network.

### 2.6. Spatial Measurements

We introduced several metrics, including network strength, network size, network center of mass, network-averaged spatial similarity, and network engagement range, to examine developmental changes in the spatial organization of brain networks. It is important to highlight that all of these metrics are derived from voxel intensities in the spatial map, which reflect the degree of engagement of each voxel in contributing to the overall network.

*Network Strength*: Network strength is defined as the average intensity of all voxels within a network, reflecting the level of engagement to the functional networks. Elevated network strength values indicate increased voxel involvement within the network. To compute network strength, we first applied a mask to identify voxels exceeding a Z-score threshold of 1.96 (p = 0.05), selecting voxels that significantly contribute to the network. Afterward, we calculated the network strength by averaging the intensities of these selected voxels, resulting in a singular value representing the network strength for each network in every scan.

*Network Size*: We quantify the spatial extent of a network using a global metric called network size to assess the shrinkage/expansion of brain networks over time. Changes in the network size within a particular network indicate an alteration in the number of voxels contributing to that network, thereby reflecting fluctuations in its size. In computing network size, we again began by applying a mask to identify voxels exceeding the threshold of Z = 1.96 (p = 0.05) within each network. Then, we counted the number of voxels meeting this criterion to quantify the size of the network.

*Network center of mass (NCM)*: We evaluate both the configuration and extent of a network comprehensively by introducing a novel metric termed the “network center of mass” (NCM). This measurement incorporates the concept of the center of mass (COM) [34], a theoretical point where the total mass of an object is considered concentrated but in fMRI is defined as the center of a cerebral activation cluster consisting of a defined number of voxels [34]. By leveraging the distance from the COM, the NCM provides insights into the spatial distribution of the network. The computation involves applying a mask to filter voxels surpassing Z = 1.96 (p = 0.05) within each network. Subsequently, we determine the COM for each network using the following formula:

Let *x*, *y*, and *z* represent the indices of the three dimensions (rows, columns, slices), and I(*x*, *y*, *z*) be the value of the voxel at (*x*, *y*, *z*).

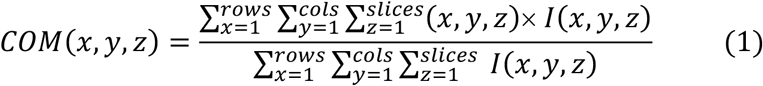

Once the COM has been obtained, we calculate the weighted average distance from COM. This involves determining the weighted distance of each voxel from the COM and subsequently calculating the average distance among these weighted distances. The formula for this process is as follows:

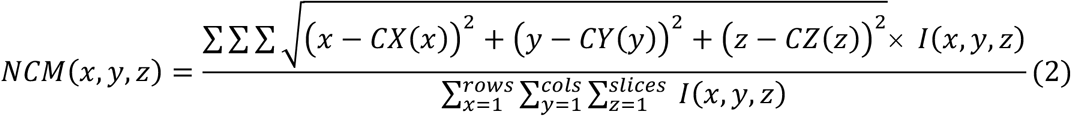

In this formula, CX(x), CY(y), and CZ(z) are the coordinates of the center of mass, I(*x*, *y*, *z*) is the value of the voxel at (*x*, *y*, *z*), and *NCM*(*x*, *y*, *z*) is the weighted average distance of all voxels within a network to COM for each network.

*Network-averaged spatial similarity (NASS)*: We introduced the “network-averaged spatial similarity” (NASS) metric to quantify how closely each participant-specific spatial map aligns with the group-level map. The group-level map represents the commonalities across all participants, reflecting the the shared spatial organization of brain networks that are consistent across individuals. To compute NASS, we calculate the correlation between each participant’s individual spatial map and the group-level network, resulting in a NASS value for each network in every scan. A higher NASS indicates that an individual’s network closely aligns with the shared spatial pattern observed across the group, suggesting that their network structure is more homogenized. Conversely, a lower NASS suggests a greater deviation from the group-level pattern, indicating more individual-specific variation in the network’s spatial map.

*Network Engagement Range (NER):* We assessed the intensity range of voxels within each network by introducing a novel metric termed “network engagement range”. This metric is used to quantify the uniformity of voxel engagement within the network. A wider intensity range may indicate greater variability in voxel contribution, which could suggest a broader spatial distribution of engagement. Conversely, a narrower intensity range might indicate reduced variability, implying more consistent voxel contribution across the network. Calculating this range involves identifying the highest and lowest voxel intensities, and the difference between these values constitutes the network engagement range, representing the variance between the maximum positive and minimum negative values of voxels within each network.

### 2.7. Statistical Analysis

For the statistical analysis, we employed a generalized additive model (GAM) [35] to explore the relationships between each metric and the variables of interest. In this model, corrected age, sex, scanner, and head motion were included as fixed effects, while individual participants were modeled as random effects. This approach allows us to account for potential linear and non-linear relationships and adjust for confounding factors, providing more robust and flexible inferences.

Effective degrees of freedom (EDF) were reported to reflect the complexity of the smoothing functions applied within the GAM. The EDF values give insight into the model’s flexibility, with higher values indicating more complex relationships captured by the model [36]. We assessed the degree of non-linearity using the EDF in our analysis, in which value of 1 indicates a linear relationship, while a value between 1 and 2 suggests a weakly non-linear relationship. Values greater than 2 signify a highly non-linear relationship, indicating that the curve is capturing more complex, non-linear patterns in the data [37]. To manage the multiple comparisons issue, p-values were adjusted using the false discovery rate (FDR) correction, ensuring that the reported significant findings are not due to random chance.

## 3. Results

Our results are divided into two main sections: Initially, we explore the functional brain networks identified in infants, conducting voxel-level analysis and modularity analysis to examine their association with age. Subsequently, we focus on the spatial developmental metrics of these networks, investigating how they evolve over time by analyzing changes in our measurements. Additionally, we compare the linear and non-linear developmental trajectories across different networks.

### 3.1. Summary of the identified brain networks and their correlations with age

We performed ICA to extract 20 components on rs-fMRI data from 74 participants and successfully identified 13 distinct brain networks in infants. These networks are presented in Fig. 1.

**Fig. 1.**
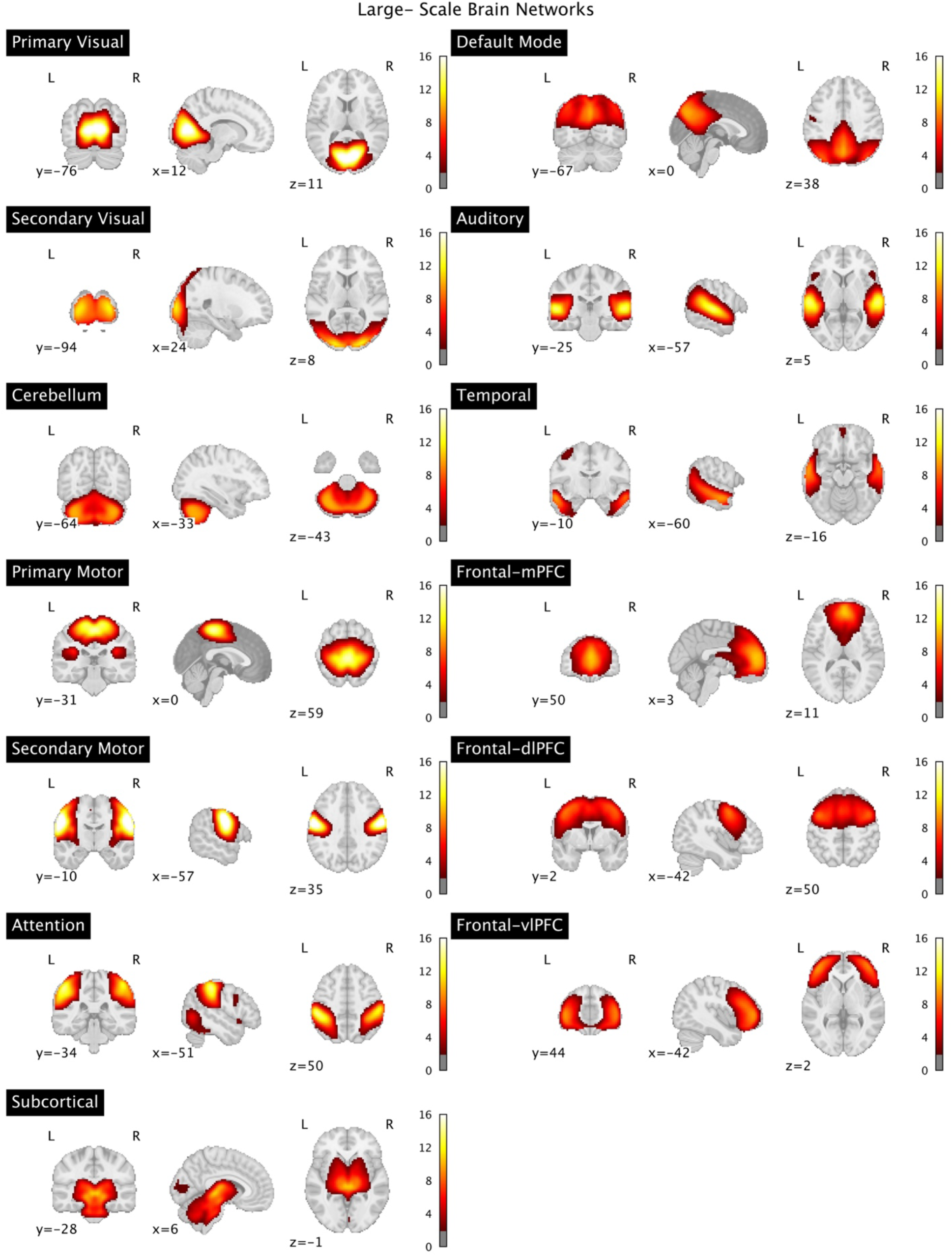
Sagittal, coronal, and axial views illustrate the z-scored voxel intensity distributions of spatial maps for 13 functional brain networks in infants.

These networks include the primary and secondary visual networks, subcortical network, cerebellum network, primary and secondary motor networks, attention network, default mode network, temporal network, auditory network, frontal-medial prefrontal cortex (mPFC) network, frontal-dorsolateral prefrontal cortex (dlPFC) and frontal-ventrolateral prefrontal cortex (vlPFC) networks. Fig. 2 illustrates the correlation between voxel intensity and corrected age. Most networks show predominantly positive correlations, suggesting that the engagement of these brain regions strengthens over the first six months of life. However, some networks also display regions of negative correlation. All results are adjusted for multiple comparisons using FDR correction.

**Fig. 2.**
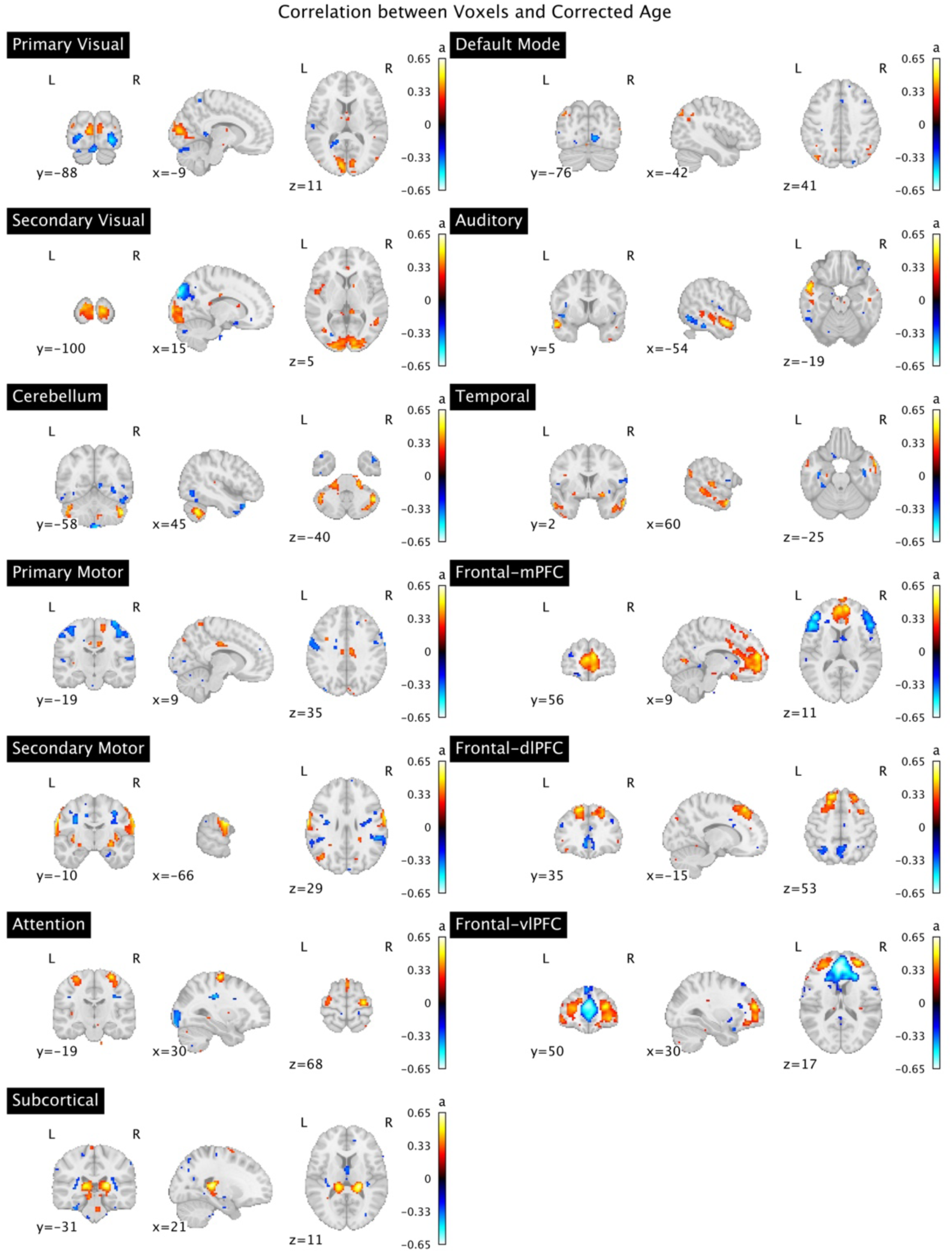
Sagittal, coronal, and axial views of the correlation between the voxel-level spatial maps of 13 functional infant brain networks and corrected age for all scans. Analyses were corrected for multiple comparisons using a 5% false discovery rate (FDR).

Another aspect of our analysis focused on examining the age-correlated voxel mean (ACVM) across networks (See 2.5 in the method for more details). Our modularity analysis on these ACVMs identified two primary modules within our networks. The first module encompasses attention, frontal-vlPFC, frontal-mPFC, frontal-dlPFC, default mode, primary motor, and secondary visual networks. The second module consists of primary visual, secondary motor, cerebellum, subcortical, auditory, and temporal networks. Fig. 3. depicts the two identified modules and shows the correlations between all networks within the first and second module.

**Fig. 3.**
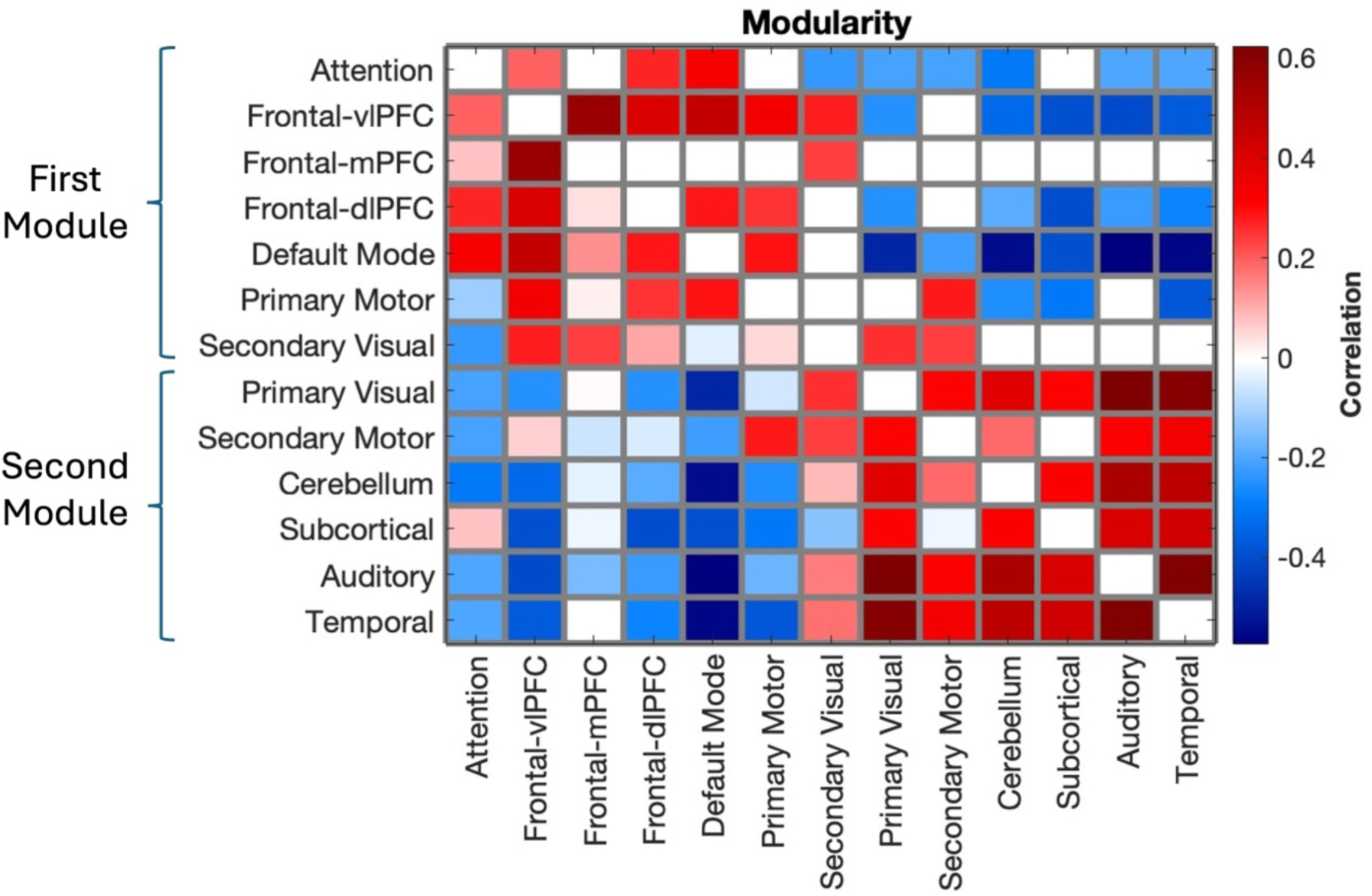
Correlation between the network belonging to the first module, comprising attention, frontal-vlPFC, frontal-mPFC, frontal- dlPFC, default mode, primary motor, and secondary visual networks, and the network belonging to the second module, including primary visual, secondary motor, cerebellum, subcortical, auditory, and temporal networks. We display only those correlation values that remained statistically significant after FDR correction for all p-values.

### 3.2. Spatial developmental metrics of brain networks

Age-related changes in spatial characteristics of all 13 networks are shown in Fig. 5 (primary visual, secondary visual, cerebellum, primary motor, secondary motor, attention, subcortical, default mode, auditory, temporal, frontal-mPFC, frontal-dlPFC, and frontal-vlPFC networks). The results reveal that network-averaged spatial similarity shows a notable increase across ages for all networks (*F* > 5.45, *p* < 0.02; Fig. 4) except for the subcortical network. Additionally, a significant decline in network engagement range is observed across most networks, including the primary visual, primary motor, secondary motor, temporal, cerebellum, attention, default mode, and frontal-vlPFC networks. However, no such decline is seen in the secondary visual, subcortical, auditory, frontal-mPFC, and frontal-dlPFC networks (*F* > 6.27, *p* < 0.02; Fig. 4) . Network strength shows a significant increase in the primary and secondary visual networks, primary and secondary motor networks, as well as in the temporal, frontal-mPFC, and frontal-vlPFC regions. Conversely, no age-related changes were observed in the remaining networks, including the cerebellum, attention, subcortical, default mode, auditory, and frontal-dlPFC networks. (*F* > 5.08, *p* < 0.04; Fig. 4).

**Fig. 4.**
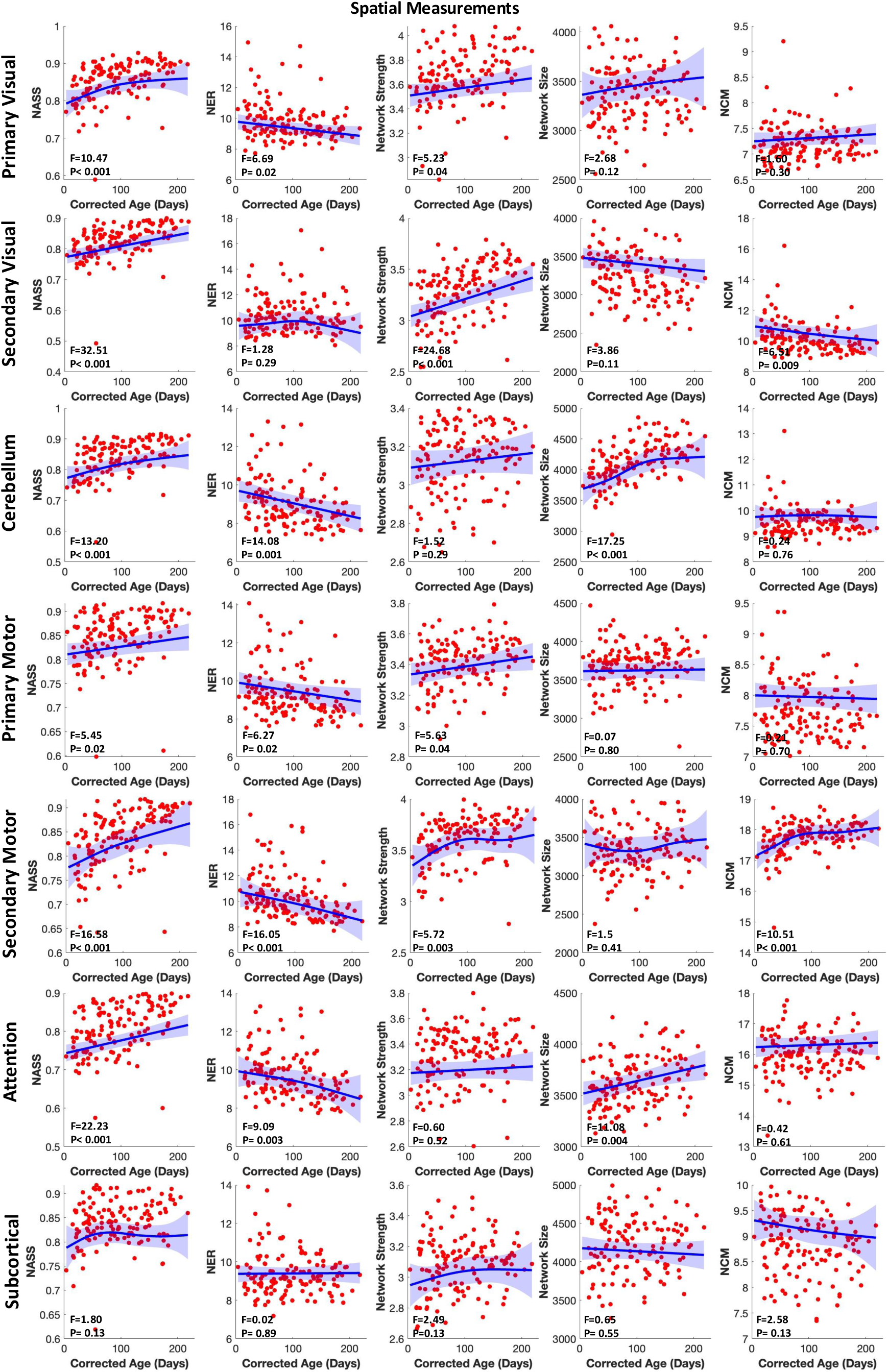
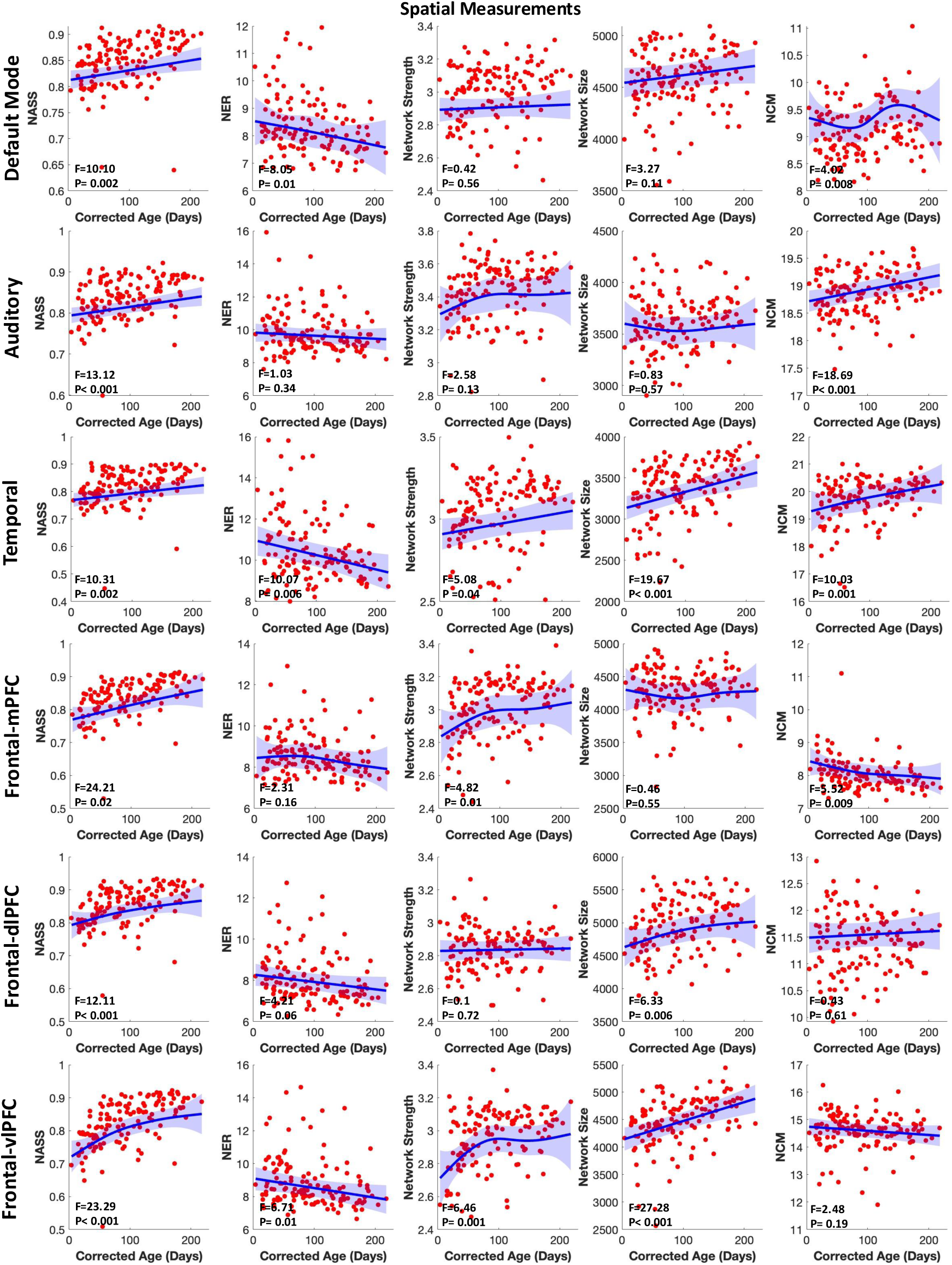
Age-related changes in spatial characteristics of the primary visual, secondary visual, cerebellum, primary motor, secondary motor, attention, subcortical, default mode, auditory, temporal, frontal-mPFC, frontal-dlPFC, and frontal-vlPFC network. NASS: network-averaged spatial similarity; NER: network engagement range; NCM: network center of mass. Each red dot represents an individual scan, while the blue line represents group mean trajectories. Shaded bands represent 95% simultaneous confidence bands. Analyses were adjusted for multiple comparisons by applying FDR correction.

**Fig. 5.**
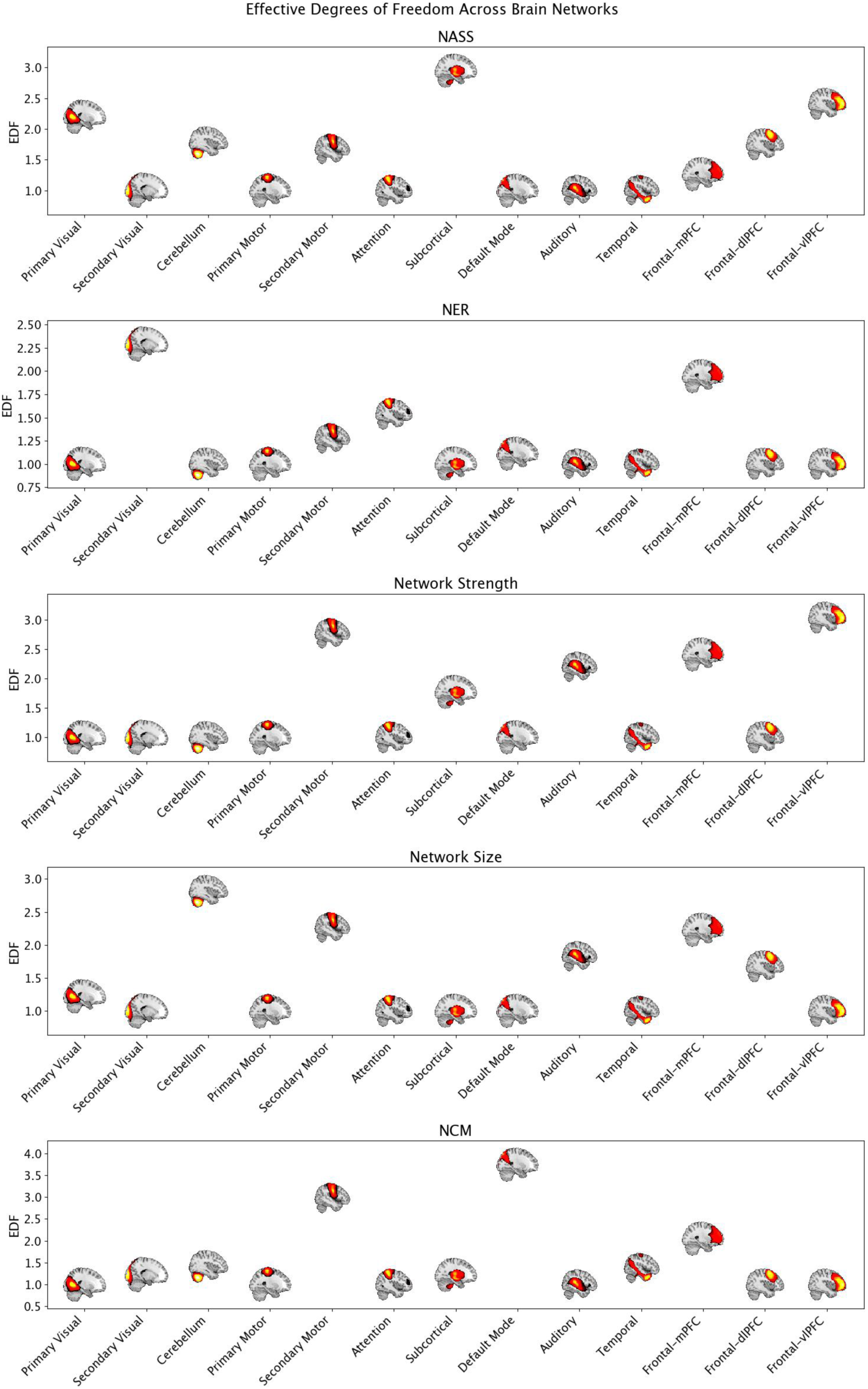
This figure displays the effective degrees of freedom (EDF) across various brain networks for different measures: NASS: network-averaged spatial similarity; NER: network engagement range; NCM: network center of mass, network strength, and network size. Each subplot represents the EDF for distinct brain networks, including the primary visual, secondary visual, cerebellum, primary motor, secondary motor, attention, subcortical, default mode, auditory, temporal, frontal-mPFC, frontal- dlPFC, and Frontal-vlPFC networks. The plots highlight variations in EDF across these networks, with each brain region’s involvement visually represented in the corresponding brain images.

Analyses of age-related change in network size reveal substantial growth in the cerebellum, attention, temporal, frontal-dlPFC, and frontal-vlPFC networks (*F* > 6.33, *p* < 0.006; Fig. 4). Lastly, network center of mass shows a significant decrease for the frontal-mPFC and secondary visual networks *F* > 0.11, *p* < 0.001; Fig. 4), while it remains unchanged for the primary visual, subcortical, cerebellum, primary motor, attention, fontal-dlPFC and frontal-vlPFC networks. In contrast, NCM experiences a significant increase in secondary motor, temporal, default mode, and auditory networks *F* > 4.02, *p* < 0.008; Fig. 4). All the p-values are after FDR-correction.

### 3.3. Comparative Analysis of Linear and Non-Linear Developmental Trajectories in Brain Networks

In this section, we compared the degree of non-linearity across networks for the five metrics: network-averaged spatial similarity, network engagement range, network strength, network size, and network center of mass. For the network-averaged spatial similarity metric, the subcortical network, along with the primary visual and frontal-vlPFC networks, exhibited the highest EDF, indicating highly non-linear trajectories. The cerebellum, secondary motor, frontal-dlPFC, and frontal-mPFC networks showed weakly non-linear trajectories, while the remaining networks had linear trajectories. In contrast, the network engagement range metric presented a different pattern. The secondary visual network was the only one with a highly non-linear trajectory. Additionally, the secondary motor, attention, default mode, and frontal-mPFC networks exhibited weakly non-linear trajectories, whereas the other networks displayed linear trajectories (Fig. 5).

For the network strength metric, the secondary motor, frontal-mPFC, frontal-vlPFC, and auditory networks exhibit highly non-linear trajectories. The subcortical network displays weak non- linearity, while the remaining networks have linear trajectories. Regarding network size, the secondary motor, cerebellum, and frontal-mPFC networks show highly non-linear patterns. Weak non-linearity is observed in the primary visual, auditory, and frontal-dlPFC networks, with the others remaining linear. The network center of mass metric highlights the secondary motor, frontal-mPFC, and default mode networks as highly non-linear, while the temporal, secondary visual, subcortical, and cerebellum networks display weak non-linearity. The remaining networks are linear (Fig. 5). Overall, across all five metrics, certain networks, like the secondary motor and frontal-mPFC, consistently show highly non-linear trajectories, while other networks like primary mortor, attention, and temporal networks remain predominantly linear, indicating varied degrees of complexity in network development.

## 4. Discussion

The investigation into the spatial development of large-scale brain networks during the initial six months of life holds significant implications for understanding early brain development and potential markers for neurodevelopmental disorders. The study, conducted with a cohort of 74 infants, utilizes ICA on rsfMRI data. Several studies have identified brain networks in infants using ICA and other fMRI methods. Fransson et al. (2007) used ICA to identify five networks in sedated infants under one year, including primary visual, somatosensory/motor, auditory, parietal, and prefrontal networks [10]. These networks are consistent with our findings, though we identified additional networks. Additionally, Damaraju et al. (2014) employed ICA with varying model orders, revealing different numbers of networks that generally aligned with those found in our study, highlighting consistency across datasets [38]. Fitzgibbon et al. (2020) identified 13 functional brain networks were identified in neonates using ICA and seed-based correlation. While many of their networks visually align with ours, they did not provide labels for direct comparison [39]. ICA leverages the multivariate relationships in the BOLD signal along with higher-order statistics to delineate distinct brain regions. This methodology allows the exploration of the developmental trajectory of 13 distinct brain networks, emphasizing spatial measurements such as network-averaged spatial similarity, network dynamic range, network strength, network size, and network center of mass.

The findings present a nuanced picture of spatial changes across various brain networks. Simultanously, the analysis of linear and non-linear development across different brain networks in infants reveals distinct and varied developmental patterns. Notably, networks such as the secondary motor and frontal-mPFC consistently exhibit pronounced non-linear trajectories across all five metrics. These findings align with earlier research by Gao et al. (2015) and Cao et al. (2017), which also documented non-linear progression in functional connectivity, highlighting the early specialization and complexity of brain network development [3,9,40]. Building on this, our analysis of spatial characteristics, particularly the observed increase in network strength during the first 6 months of life suggests enhanced network engagement, which may lead to greater neural activity and maturation. During this period, myelination enhances signal transmission speed and efficiency, while synaptogenesis increases the formation of new synapses, establishing essential neural circuits [41]. Concurrently, synaptic pruning refines these circuits by eliminating excess synapses, enhancing the functionality of crucial connections [41]. The establishment of neural connections integrates sensory, motor, and cognitive functions across different brain regions. These processes, coupled with increased metabolic activity to support the growing brain, contribute to the heightened voxel intensity seen in fMRI scans, reflecting the rapid and intricate maturation of functional brain networks during early development [9,41,42]. In addition, the heightened network strength could be indicative of increased contribution to the network’s function and neural activity within these brain networks may suggest a crucial developmental phase characterized by the refinement and optimization of neural circuits [43,44]. This phenomenon aligns with existing literature on early brain development, emphasizing the critical role of the first months of life in shaping the foundation for later cognitive and functional abilities [45]. Moreover, the observed patterns may serve as valuable markers for tracking neurodevelopmental trajectories and identifying potential deviations from typical developmental patterns.

The second metric we examined was network size, with a focus on its varying trajectories across different brain networks. The unique spatial dynamics observed suggest distinct developmental pathways within individual neural networks during early life. Notably, networks showing a significant increase in size may indicate regions undergoing rapid growth, likely driven by key developmental processes such as synaptogenesis [46–48]. For example, the observed increase in cerebellar network size (0.08% per day) could be attributed to the cerebellum’s rapid growth in both volume and connectivity during early development [49]. This expansion facilitates its integration into larger brain networks involved in motor and cognitive functions. Studies have shown that the cerebellum experiences significant growth during the first year of life, largely driven by processes such as synaptogenesis and the formation of new neural connections [49]. Moreover, Fossella et al. (2003) suggest that increased synaptogenesis during infancy and early childhood plays a key role in the maturation of attention networks, which are critical for cognitive functions like working memory and problem-solving [50]. This biological process is closely tied to the observed developmental expansion of network size, particularly in the attention network (0.04% per day). The growth and refinement of these networks underscore the importance of synaptogenesis in early brain development, as it facilitates the formation of essential neural circuits that support cognitive development [50]. Additionally, the observed increase in the size of the temporal (0.08%/day), frontal-dlPFC (0.06%/day), and frontal-vlPFC (0.08%/day) networks indicates significant developmental changes that could be related to the enhancing of cognitive and social functions in infants. For instance, one previous study showed, during the first two years of life, the brain’s network topology evolves, with an increase in long-distance connections and strengthening of functional connectivity, particularly in the temporal regions [51]. On the other hand, networks maintaining a static network size might indicate regions achieving structural stability or undergoing subtle, yet functionally relevant modifications not reflected in voxel quantity. This spatial heterogeneity underscores the complexity of early neurodevelopment and the necessity for a network-specific exploration to unravel diverse patterns of growth and maturation across different infant brain regions.

The noted rise in network-averaged spatial similarity, as measured by the correlation between individual spatial maps of each brain network and the group-level spatial map of the same network, reveals intriguing insights into the consistency of network organization. This phenomenon suggests a convergence towards a more standardized spatial configuration across various brain networks. The heightened NASS implies that the inherent patterns and structures characterizing each network tend to align more closely with the group-level representation over time. As the group-level map reflects the common spatial organization of brain networks shared across all participants, an increase in NASS with age indicates that an individual’s network is becoming more aligned with the group’s spatial pattern. This suggests that the individual’s network structure is becoming more homogenized over time. In contrast, a lower NASS value implies a greater divergence from the group-level pattern, indicating more individual-specific variability in the network’s spatial map. Our findings of increasing network-averaged spatial similarity align with previous studies that show an increase in spatial similarity of individual variability maps in the prenatal brain as it develops with age [52]. This trend underscores the emergence of individual variability in the brain’s functional architecture during prenatal development, potentially establishing the neural foundations that contribute to individual behavioral differences later in life [52]. In addition, NASS can reveal how network organization changes over time. Increasing NASS between the individual and group-level maps might suggest that network organization becomes more consistent with age. Moreover, it’s important to note that smoothing can influence NASS. While it increases the signal-to-noise ratio [53,54], enhancing the detection of consistent patterns across participants, it can also affect inter-participant variability by making individual differences less apparent [55]. To mitigate this effect, we carefully selected our smoothing kernel size based on the optimal range identified in prior studies for group-level ICA [56], ensuring that the benefits of smoothing are realized while minimizing its impact on inter-participant variability.

In our examination of infant brain development, we analyzed the network center of mass across various brain networks. Our results revealed a dynamic pattern marked by fluctuations in this measure: while some networks showed an increase (such as the secondary motor, temporal, and auditory networks), others exhibited a decrease (including the secondary visual and frontal networks), while certain networks remained stable. This variability highlights the complex nature of brain network development during early infancy. These observations regarding changes in the spatial distribution of networks align with existing literature on functional connectivity metrics, which emphasize the direct influence of physical distance between nodes on connectivity within any network [57]. In our analysis, we observed that spatial distribution changes across age, with our metric relying on the distance between the center of mass of each network and the voxels participating in that network. Similar to functional connectivity, which is known to change as a function of distance [57], the degree of voxel engagement in the functional network could also be influenced by this distance. As brain networks evolve over time, it is plausible that the spatial distance between the center of mass and participating voxels plays a role in shaping network engagement, reflecting how proximity can affect both structural and functional connectivity.

Network engagement range is another novel method that we studied. The NER quantifies the difference between the highest positive and lowest negative values of brain voxels within a specific network. A larger NER signifies a greater disparity in voxel intensity. Put simply, if the NER decreases across age, it suggests network consolidation over time, as voxel intensities become closer, indicating a reduction in random intensity fluctuations and more consistent voxel contribution across the network. In a related study, the authors utilized a measure called gradient range instead of the NER [58]. While the gradient range differs from our NER and this difference arises because they computed the gradient range instead of the engagement range, but their findings show an increase over time, indicating progressively maturing global topographic profiles in children. This finding aligns well with our results, supporting the notion of network consolidation during development [58]. On the other hand, the NER of the auditory network remains stable during the first six months of life (see Section 3.2 for more details). This stability may be due to the fact that the auditory system is one of the earliest functions to develop in utero. The auditory network is already functioning at birth and is among the first networks to reach an adult-like state. This early maturation enables infants to process sound shortly after birth, which may explain why no significant changes are observed in the auditory network’s NER during this early developmental period [59]. However, most of the other networks are still consolidating and going to change over these seven months, for example, Lin et al. [60] have shown that the strength and extent of sensorimotor networks increase more rapidly than those of visual networks throughout early childhood, that is consistent with our results because not only the NER decreasing in primary and secondary motor networks, but also the rate of decreasing is much more than visual networks. Fluctuations in the network engagement range across diverse age groups may signify alterations in the range of brain activity. Moreover, changes in the NER may indicate reorganization within brain networks. A decreasing range might indicate more consistent and predictable contribution to the network, while an increasing range could suggest greater variability within the network. When examining the linear and non-linear developmental trajectories of spatial characteristics discussed earlier, the non-linear patterns observed in networks such as the subcortical network, especially when considering metrics like network-average spatial similarity, network center of mass, and network strength, align with findings from Alex et al. (2023). Their research demonstrated that subcortical structures, such as the amygdala and thalamus, undergo intricate and extended maturation processes during infancy and early childhood [61]. These observations suggest varied spatial growth patterns, indicative of the complex interplay between neural growth and pruning during early developmental stages [40]. One study indicated that brain myelination occurs rapidly within the first six months of life, but this process is not uniform across different brain regions [62]. Areas associated with vision and motor functions, particularly in the posterior and superior parts of the brain, experience faster myelination, aligning with the critical development of these abilities during infancy. This accelerated myelination may explain the non-linear trajectories observed in visual and motor networks in our findings. Overall, these results underscore the diverse and non-linear development of brain networks in infants, highlighting the crucial role of early specialization and integration in shaping cognitive and sensory functions [52]. Lastly, further research is warranted to delve deeper into the specific mechanisms contributing to these observed changes in network strength, network size, NCM, NASS, and NER to explore their potential implications for long-term neurodevelopmental outcomes. When considering all these observations together, the conclusion is that the early months of life are marked by a critical phase of brain network refinement and optimization. The increasing spatial similarity within networks, coupled with changes in network engagement range and size, indicates that the brain is not only becoming more organized but also more functionally specialized. These developments are essential for laying the groundwork for later cognitive and behavioral abilities. The consistency in findings across different metrics further supports the idea that early neurodevelopment is characterized by both the consolidation of key networks and the ongoing reorganization of others to meet the growing demands of a developing brain.

In age-associated voxel-level analysis, we observed predominantly positive correlations between voxel intensity and corrected age in several key brain networks, including the visual, motor, and auditory regions. This indicates that these networks are strengthening during the first six months of life, which aligns with their crucial role in the early development of sensory and motor functions [9,43,45]. Conversely, some negative associations were found in the vlPFC-frontal, dlPFC-frontal, and mPFC-frontal, regions known for their involvement in higher-order cognitive functions. These negative correlations may reflect early synaptic pruning or the ongoing refinement and specialization of these prefrontal regions, which continue to develop throughout childhood and adolescence [63–65]. After that the modularity analysis identified two primary modules within the brain networks of infants, reflecting distinct patterns of network interactions. The first module, encompassing attention, frontal-vlPFC, frontal-mPFC, frontal-dlPFC, defult mode, primary motor, and secondary visual networks, aligns with regions involved in higher-order cognitive functions and integrative processes. This module’s composition suggests a coordinated development of cognitive control, attention regulation, and sensory processing during early infancy [66,67]. The second module, consisting of primary visual, secondary motor, cerebellum, subcortical, auditory, and temporal networks, indicates a focus on sensory processing, motor control, and basic cognitive functions. One study examining the functional connectivity of rs-fMRI in infants aged 0-1 year provides valuable insights into how brain functional networks develop during this critical period. The research reveals that as infants grow, the brain’s functional network becomes increasingly subdivided into a greater number of distinct functional modules [68]. This finding aligns with our study’s observations, which identify a division of brain networks into two primary categories: higher-order cognitive functions and sensory processing/motor control. The gradual subdivision of functional modules supports the notion that, even from an early age, the brain organizes itself into specialized networks that cater to different cognitive and sensory-motor functions. The segregation into two distinct modules highlights the early specialization and integration of functional brain networks, essential for developing complex behaviors and cognitive abilities in infants [9,69].

## Limitations

This study offers valuable insights into the normative development of brain networks during the crucial first six months of life. However, there are several limitations to consider. A more pertinent limitation in this study, which is explicitly designed to map neurotypical developmental trajectories, is the lack of diversity in terms of race, ethnicity, and socioeconomic status. These factors could have effects in shaping brain development due to their influence on environmental exposures, healthcare access, and social determinants of health. As such, the limited diversity in the sample may restrict the generalizability of the findings, as the developmental trajectories observed may not fully represent the experiences of infants from more diverse backgrounds. Additionally, while the sample size provides preliminary insights, it may not be sufficient to generalize these findings across the entire infant population. Moreover, the study predominantly emphasizes spatial changes in brain networks, yet it does not deeply explore the temporal dynamics that are essential to understanding how these networks evolve over time and their role in the developmental trajectory.

## Future Work

Future research should incorporat a more diverse cohort of infants, particularly those at risk for neurodevelopmental disorders, to gain a comprehensive understanding of how deviations from typical development manifest in brain network organization. A crucial area for future exploration is the study of the presence and development of 105 Intrinsic Connectivity Networks (ICNs) derived from brain imaging data, representing different levels of brain connectivity across multiple spatial scales. These templates capture both large-scale global networks and smaller, more localized brain circuits, providing a comprehensive view of functional connectivity in the brain multiscale ICNs templates in infants. Understanding how these ICNs emerge and evolve during early development could provide critical insights into the foundational architecture of brain networks. Investigating how these spatial maps of brain networks change over varying time scales from short-term fluctuations to long-term changes could offer valuable insights into the neural processes underlying cognitive and behavioral development [13]. This approach could also help identify early markers of neurodevelopmental disorders, facilitating earlier and more targeted interventions.

## Conclusion

In conclusion, this study provides a comprehensive examination of the spatial development of large-scale brain networks during the first six months of life, offering critical insights into the normative processes of early brain maturation. By leveraging advanced metrics such as network strength, size, center of mass, spatial similarity, and engagement range, we have delineated the intricate patterns of neural development that underscore the foundation of cognitive and sensory functions. Our findings highlight the complexity and heterogeneity of early brain development, characterized by both the consolidation of key networks and the ongoing reorganization of others. These early months are pivotal for laying the groundwork for later cognitive and behavioral abilities, with distinct spatial dynamics within individual networks potentially serving as markers for typical and atypical neurodevelopmental trajectories. While our study emphasizes normative development, future research should expand to include at-risk populations and longitudinal studies to further explore the implications of these early spatial patterns for long-term outcomes. This research not only contributes to the understanding of early brain development but also holds potential for early detection and intervention strategies in neurodevelopmental disorders.

