## Supplemental Material for "Spatial Development of Brain Networks During The First Six Postnatal Months"

*1.1. Quality Control*

We conducted quality control (QC) on the preprocessed fMRI data using the NeuroMark framework [1]. Scans that did not show good normalization to the MNI standard space were excluded from further analysis. We compared individual masks with the group mask and retained subjects with a high similarity between the two. Initially, based on the first fMRI time volume, we created individual masks for each subject by setting voxels to 1 if they exceeded 90% of the whole brain mean. We then generated a group mask by setting voxels to 1 if more than 90% of subjects had a value of 1 in their individual masks. For each subject, we calculated spatial correlations between the group and individual masks using voxels within the top 10 slices, bottom 10 slices, and the entire mask, resulting in three correlation values per subject. Subjects were included for further analysis if their top-10-slice correlation was greater than 0.75, bottom-10-slice correlation was greater than 0.55, and whole-brain correlation was greater than 0.8. This method ensures high-quality masks and fMRI data, a technique proven effective in our previous studies. Due to the requirements of the NeuroMark [1], scans with fewer than 100 time points were excluded. The average scan duration was 5 minutes and 54 seconds. Ultimately, 74 subjects (137 scans) met all QC criteria and were included in the analysis.

*1.2. Figures*

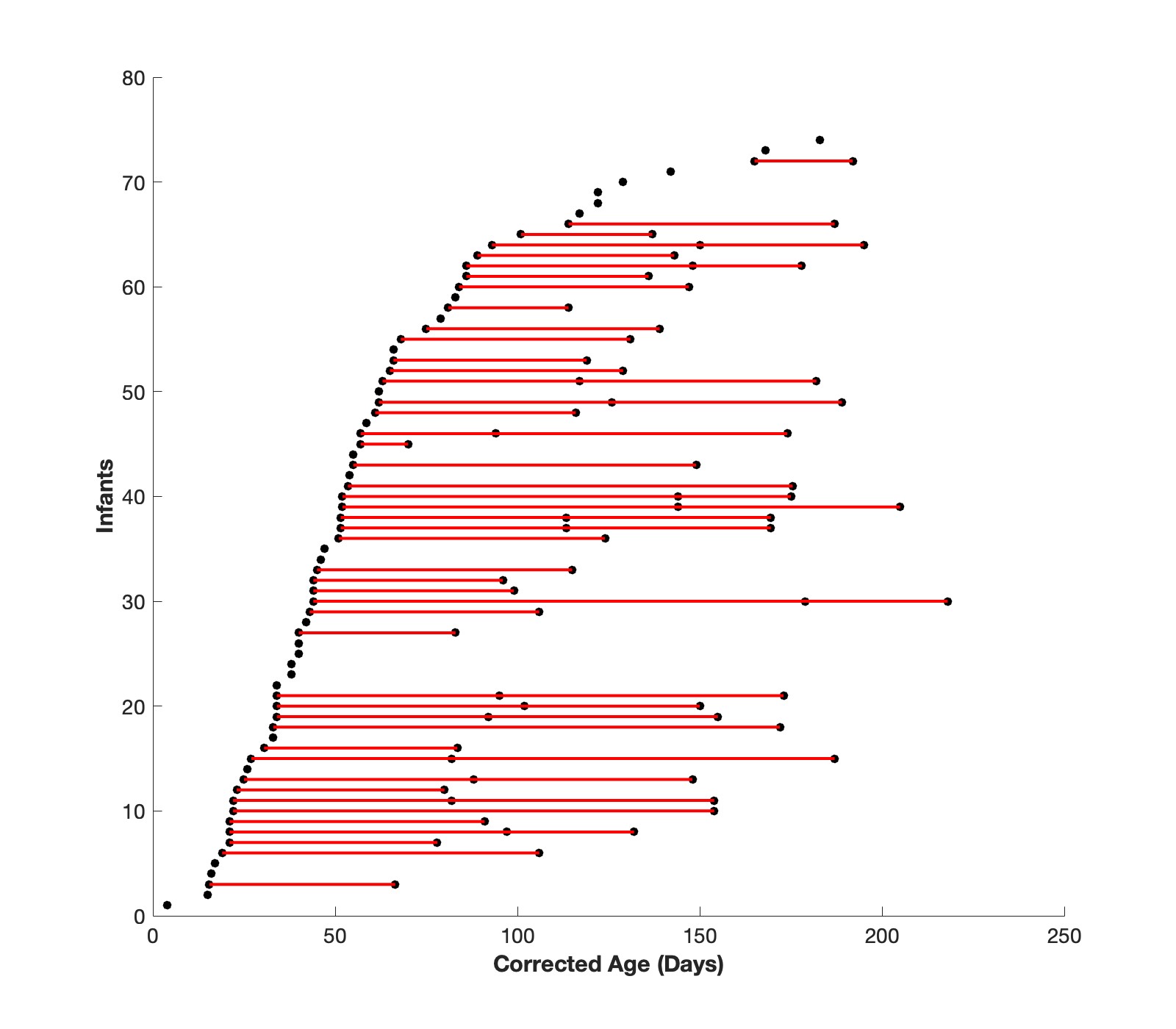

Fig. S1 The distribution of scans by corrected age for all included participants. Each dot represents one scan from a participant and dots connected by lines represent all included longitudinal scans from that given participant. Data were collected using a non-uniform longitudinal sampling design, with scans collected from each infant at up to 3 pseudorandom time points between birth and 6 months (137 scans total, each separated by a mean of 1.4 days (SD = 1.8)

*2. References*

[1]	Du Y, Fu Z, Sui J, Gao S, Xing Y, Lin D, et al. NeuroMark: An automated and adaptive ICA based pipeline to identify reproducible fMRI markers of brain disorders. Neuroimage Clin 2020;28. https://doi.org/10.1016/J.NICL.2020.102375.
